# Disparate social structures are underpinned by distinct social rules across a primate radiation

**DOI:** 10.1101/2025.07.31.667759

**Authors:** Jacob A. Feder, Susan C. Alberts, Elizabeth A. Archie, Małgorzata E. Arlet, Alice Baniel, Jacinta C. Beehner, Thore J. Bergman, Alecia J. Carter, Marie J. E. Charpentier, Kenneth L. Chiou, Catherine Crockford, Guy Cowlishaw, Federica Dal Pesco, David Fernández, Julia Fischer, James P. Higham, Elise Huchard, Auriane Le Floch, Julia Lehmann, Amy Lu, Gráinne M. McCabe, Alexander Mielke, Liza R. Moscovice, Benjamin Mubemba, Megan Petersdorf, Caroline Ross, India A. Schneider-Crease, Robert M. Seyfarth, Noah Snyder-Mackler, Larissa Swedell, Franziska Trede, Jenny Tung, Anna H. Weyher, Roman M. Wittig, Jason M. Kamilar, Joan B. Silk

## Abstract

Over six decades of research on wild baboons and their close relatives (collectively, the African papionins) have uncovered substantial variation in their behavior and social systems. While most papionins form discrete social groups (single-level societies), a few others form small social units that are nested within larger supergroups (multi-level societies). These two systems are generally thought to be qualitatively distinct, but data from wild populations increasingly suggest that there may be areas of overlap. To quantify this potential gradient in social structure, a more systematic, comparative analysis is needed. Here, we constructed a database of behavioral and demographic records spanning 135 group-years, 28 social groups, 13 long-term field studies, and 11 species to quantify variation in grooming network structure, and identify the individual and dyadic properties (e.g., kinship and social status effects) that underlie this variation. Consistent with accumulating observations in the field, the single-level species could be divided into two categories: *cohesive* and *cliquish*. Cohesive single-level networks were dense, kin-biased, and moderately rank-structured, while cliquish single-level networks were more differentiated, slightly more kin-biased, and strongly rank-structured. As expected, multi-level networks were very modular and shaped by females’ ties to specific dominant males but varied in their kin biases. Taken together, these data suggest that (i) kin and rank biases are widespread but vary in their strength; (ii) male-centered subgroups are exclusive to multi-level systems; and (iii) increases in network modularity can emerge in response to heightened nepotism and male-centered clustering.

**SIGNIFICANCE STATEMENT:** What forces explain variation in primate societies? While kinship and dominance shape the social lives of many of our close relatives, it is unclear how their effects differ across species. Using a new database comprising decades of field research, we found that baboons and their close relatives fell into three general patterns: one in which groups were cohesive, kin-biased, and moderately rank-biased, another in which groups were more cliquish and nepotistic, and a third in which groups were divided into clusters centered on dominant males. Distinct primate societies may thus reflect differences in the strength of females’ nepotistic biases and the degree of males’ social influence.

## INTRODUCTION

Behavioral ecologists have long sought to identify the selective pressures that have shaped the evolution of sociality in mammals (1–3) and primates in particular (4–6). Theoretical and empirical work suggests that extrinsic factors, such as the distribution of food and the intensity of predation risk (7–10), shape the size, composition, and cohesion of social groups (i.e., social organization: 11), while intrinsic factors, such as kinship, reciprocity, and local competition (12–15), influence the tenor, frequency, and stability of interactions within groups (i.e., social structure: 11). On the whole, these theoretical frameworks have been moderately successful in linking ecological factors with the size and composition of social groups (16, 17), but markedly less successful in predicting the emergence of dominance hierarchies, kin structures, and differentiated relationships within groups (reviewed in 18, 19). This shortcoming likely stems from the historical lack of fine-grained data necessary to quantify variation in social network structure. However, longitudinal, individual-based field studies (20), collaborative science initiatives (21), and social network methods (22) have increasingly facilitated the construction of comparative social interaction databases (e.g., MacaqueNet: 23, DomArchive: 24, Animal Social Network Repository: 25, reviewed in 26).

Here, we compiled a new database of behavioral and demographic data from 13 long-term field studies of baboons and their close relatives: collectively, the African papionins (**Figure 1**). The Comparative Analysis of Papionin Societies (CAPS) database is collaborative, standardized, and designed to explore links between ecology, demography, and social structure. The papionins are an ideal lineage for assessing the processes that shape social structure, as they encompass substantial variation in their ecologies, dietary niches, and social systems (reviewed in 27). While most papionin species form single-level societies in which individuals belong to one coherent social group (i.e., the normative social system observed in group-living primates), a few species form multi-level societies in which individuals belong to small, one-male units that are nested within higher, more inclusive social tiers (28). Multi-level societies have evolved at least twice in the papionins (29), once in geladas (*Theropithecus gelada)* and once or twice in baboons (presumably in the common ancestor of *Papio hamadryas* and *P. papio*). Moreover, the papionins have been studied extensively over the last 60 years, owing in large part to their use as model systems in human evolutionary research (30–34).

**Figure 1.**
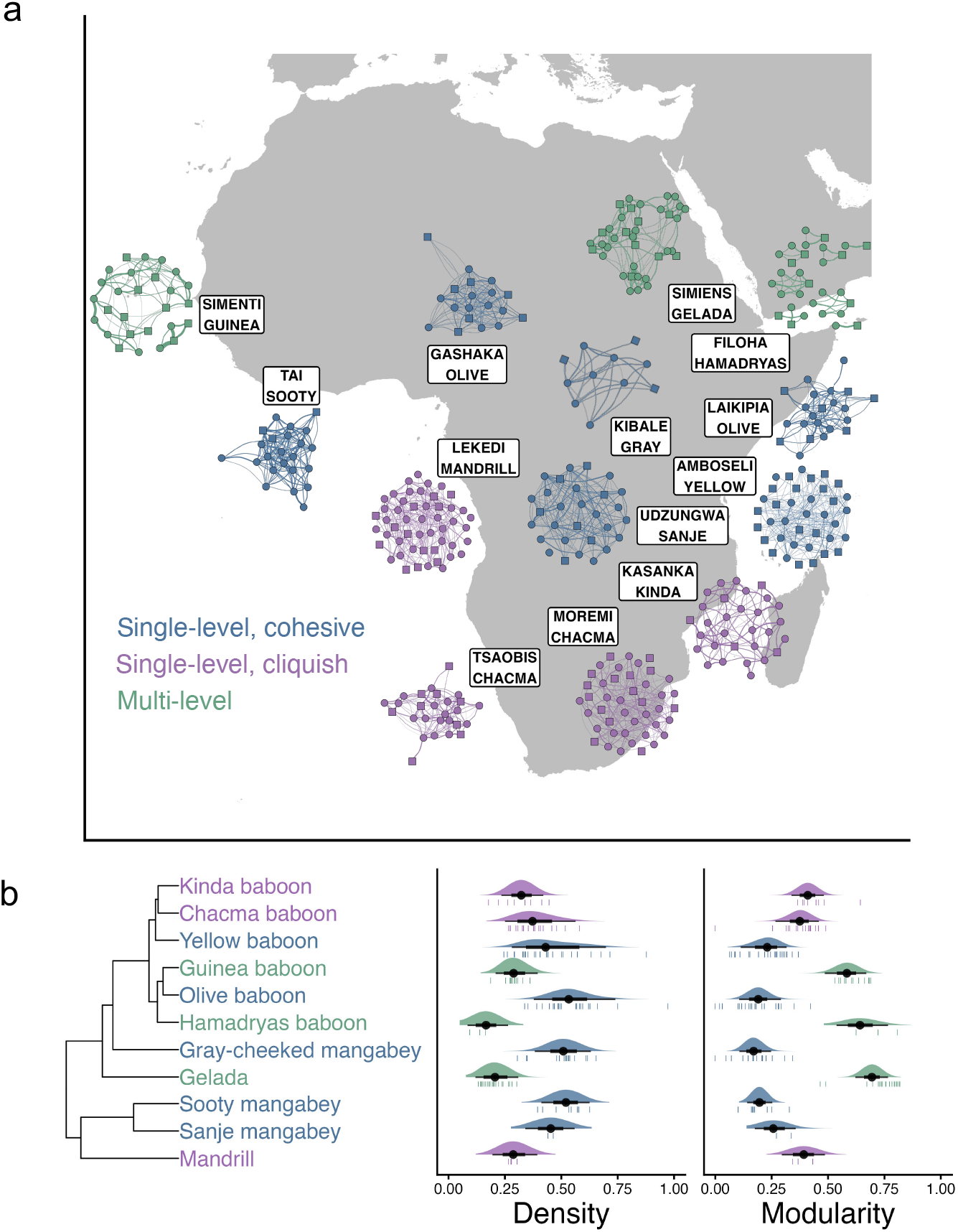
Grooming networks varied across the papionins. (a) Representative grooming networks across the papionins, provided over a map of collaborating field research sites. Within each network, circles indicate females, and squares indicate males. The width of each edge corresponds to the proportion of time or scans each dyad was observed grooming or the rate of grooming interactions (undirected, to improve visibility). (b) The density and modularity of each grooming network within the sample, organized by species, and provided alongside a phylogeny of the papionins, from Kuderna et al. (70). Species-level predicted density distributions were generated from the models described in the Main Text, including both fixed and random effects. Thick bars indicate 66% probability intervals, while thin bars indicate 95% probability intervals. Tick marks below each interval indicate observed network metrics for each group-year. In (a) and (b), each network and network measure are colored according to the social categories described in the Main Text (blue = cohesive, single-level; purple = cliquish, single-level; green = multi-level).

The single-level (mangabeys, mandrills, and most baboon species) and multi-level papionins (Guinea baboons, hamadryas baboons, and geladas) show species-specific differences in their group sizes, dispersal patterns, competitive regimes, and social strategies (27, 35–37). In the single-level societies, females typically remain in their natal groups, develop strong relationships with their maternal kin, and form matrilineal dominance hierarchies in which related females typically occupy adjacent ranks (38–42). In multi-level geladas, females similarly remain in their natal units, form matrilineal hierarchies, and develop strong kin bonds (43, 44), mirroring the patterns found in single-level groups. By contrast, in multi-level hamadryas and Guinea baboons, females leave their natal units (45, 46), limiting opportunities for females to form lasting ties with maternal relatives. Female dispersal is voluntary in Guinea baboons but not in hamadryas baboons, in which females are forcibly transferred between units during male ‘takeover’ events (47). Strong female-male relationships are also observed in both single-level (48–51) and multi-level papionin societies (52–54). However, the patterning of these relationships varies across the papionins, likely due to differences in the intensity of male-male mating competition (55), the degree of female mate choice (49, 52), and the benefits individuals might derive from forming durable opposite-sex bonds (15, 56).

While single-level and multi-level societies have historically been treated as separate, discrete phenomena (29), there may be areas of overlap between these two systems. For instance, chacma baboon groups can fracture into temporary one-male subgroups – a pattern that has been likened to the formation of one-male reproductive units within multi-level societies (57, 58, reviewed in 59).

Kinda baboons form durable opposite-sex associations that superficially resemble those found in multi-level units (49), and mandrills can form very large and putatively clustered groups (60, but see 61). Variation within the single-level societies may thus provide insights into how and why some lineages began forming larger and more modular groups. Standardized behavioral data are needed to place the single-level and multi-level societies along such a potential continuum, identify the generative ‘rules’ (e.g., kin and rank biases, popular male effects) that produce variation in social structure across this clade, and point to the potential evolutionary drivers of distinct social strategies.

In this inaugural study, we construct social networks using grooming interactions, which have long been used to quantify the strength and tenor of primate social bonds (62). Using these networks, we (i) assess variation in grooming network structure (e.g., density and modularity) across the papionins and evaluate its consistency with qualitative categories; (ii) measure the strength of nepotism and shared male preferences on female-female grooming relationships; and (iii) quantify the impact of both male and female dominance rank on female-male grooming relationships. Taken together, these analyses provide a more comprehensive synthesis of papionin social systems, capturing variation both within and between the single- and multi-level societies. In constructing the CAPS database, we also lay the groundwork for future research targeting the ecological pressures and evolutionary forces that shape social structure in primates and social animals more broadly.

## RESULTS

The CAPS database includes grooming data from 11 species across 13 sites, representing a total of 135 group-years of behavioral data from 28 distinct social groups (group-years per social group: mean = 4.8; range = 1-11; **Table S1**). While behavioral sampling typically lasted throughout the calendar year, observations were occasionally restricted to particular seasons (see *Supporting information*). When constructing annual networks for the multi-level societies, we aggregated data at the party-level for Guinea baboons, the clan-level for hamadryas baboons, and the band-level for geladas. While the apt unit for comparison across evolutionarily distinct social systems is not always clear (63), these three groupings reflect the first general level above the one-male unit and capture their tiered network structures. The papionin networks varied widely in their size and composition (3-58 females, 1-20 males, 0.75-12.0 females-per-male; **Table S1**). In general, female-female grooming was more common than female-male or male-male grooming. Within female-male dyads, grooming from females to males typically exceeded grooming from males to females, except in Kinda baboons (**Figure S1**). Male-male grooming occurred regularly in Guinea baboons and geladas (**Figure S1**) but was uncommon in the single-level societies. While grooming is common between and among solitary and follower hamadryas males (64), these extra-unit interactions were not captured within the hamadryas dataset (65).

We constructed weighted, directed grooming networks for each group-year within the dataset (n = 135, see **Figure 1** for representative networks), including all adult and subadult individuals. Then, we constructed a Bayesian edge model for each group-year within the dataset using ‘bisonR’ (66), which captured network edge weights as (i) the proportion of observation time that dyads were observed grooming or (ii) the rate of grooming events, accounting for differences in sampling effort. Because these two sampling types led to distinct edge weight definitions that are not wholly equivalent, we divided each edge weight by the mean value observed in its respective group-year to standardize data within group-years and facilitate relative comparisons (67). By including observation effort as an ‘exposure’ term, we also accounted for uncertainty when generating edge weights, assigning narrower credible intervals to dyads with greater sampling effort. After calculating edge weights and building posterior networks for each group-year, we derived two network-level metrics: *density*, which is the proportion of all dyads that were observed grooming; and *modularity*, which reflects the extent to which a grooming network can be divided into discrete clusters via a random walk process (68). Because ‘bisonR’ models assume non-zero weights for all edges, we set dyads whose edge weights fell below the minimum empirically observed value for that group-year to 0 before calculating these global network metrics (following 69).

### Papionin grooming networks vary as a function of network size and social system

Based on prior research, networks formed by multi-level species are expected to be less dense and more modular than networks formed by single-level species. However, prior researchers (49, 59, 61) have also hypothesized that, within the single-level societies, Kinda baboons, chacma baboons, and mandrills form more modular groups than yellow baboons, olive baboons, and the three sampled mangabey species. Thus, there may be more granularity in social structure than that captured by conventional single-level and multi-level categories. To quantify differences between the single-level and multi-level societies and test for substructure within the single-level species, we constructed a three-level categorical variable that divided the single-level societies into two *a priori* social categories: *cohesive* and *cliquish*. The cohesive category included olive baboons, yellow baboons, and both genera of mangabeys, while the cliquish category included Kinda baboons, chacma baboons, and mandrills. The third category included the three species that form multi-level societies (i.e., Guinea and hamadryas baboons, geladas).

To quantify the effects of these three social categories on network density and modularity, we constructed two phylogenetic regression models using ‘brms’ (71) in ‘R’ version 4.4.0 (72), which treated density and modularity as zero-one-inflated beta outcomes. Across both models, we included social system type (i.e., cohesive single-level, cliquish single-level, and multi-level) and network size (i.e., the number of adults and subadults) as covariates, as global network metrics often covary with group size both within (73, 74) and across species (75, 76). This approach enabled us to determine whether categorical differences in network structure persisted after accounting for differences in network size. To account for repeated measures across biological and social levels, we controlled for species, population, group identity, and phylogenetic relatedness (using a pruned species-level phylogeny from (70): **Figure 1b**) using random effects.

In general, smaller networks were denser than larger networks (β_GroupSize_ = -0.61, 89% CI = [-0.74, - 0.49], **Figure 2a**), and both cohesive and cliquish networks were denser than multi-level networks (β_Multi-level_ = -0.99, 89% CI = [-1.64, -0.32], **Figure 2b**). However, networks of species categorized as cohesive and cliquish were equally dense (β_Cliquish_ = -0.32, 89% CI = [-0.98, 0.35]). Larger grooming networks were also more modular than smaller grooming networks (β_GroupSize_ = 0.33, 89% CI = [0.18, 0.51], **Figure 2c**), and species categorized as cohesive and cliquish varied in their modularity. Cohesive societies were the least modular, cliquish societies were somewhat more modular (β_Cliquish_ = 0.70, 89% CI = [0.22, 1.12]), and multi-level societies were the most modular (β_Multi-level_ = 1.83, 89% CI = [1.37, 2.26], **Figure 2d**). Thus, cliquish species formed more modular grooming networks than cohesive species, even after accounting for differences in network size. To validate this three-factor categorization, we fit naive Bayesian classifier (77) models trained on half of the dataset, which correctly recovered our assigned cohesive and cliquish social categories for 91.3% of group-years (range: 88-96% accuracy across the 20 posterior networks). By contrast, classification models trained on randomly assigned categories were correct only 60.4% of the time (range: 51.0-66.7% accuracy). Thus, these categories appear to be an apt reflection of the variation within our sample, and we used this three-level factor in all subsequent analyses.

**Figure 2.**
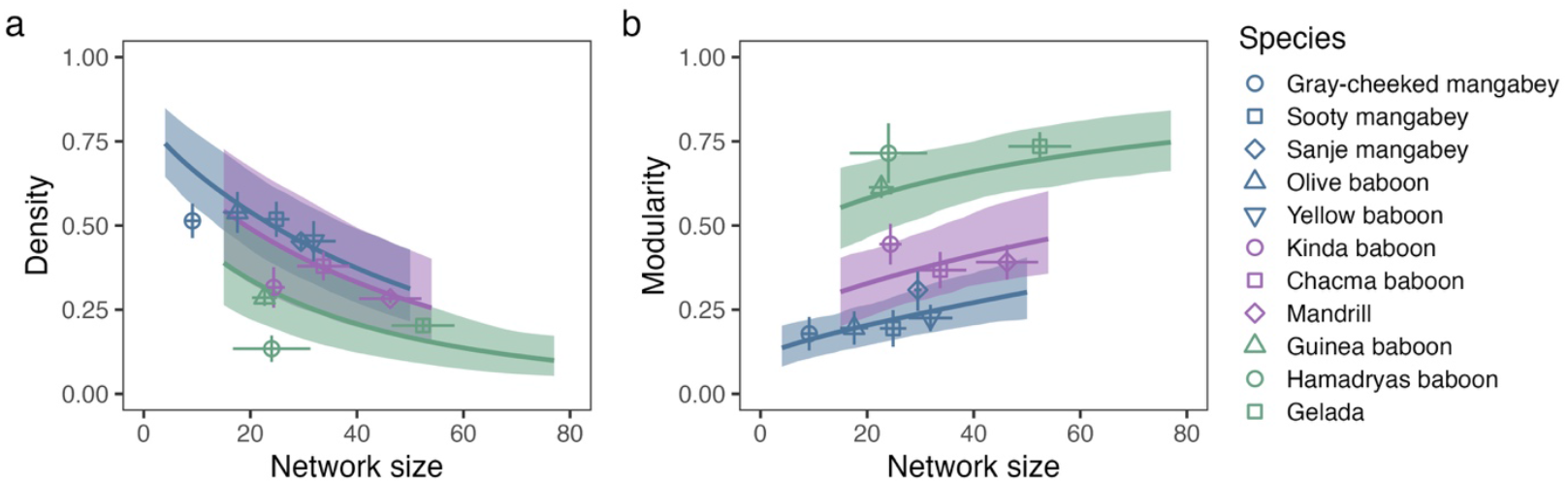
Network metrics varied as a function of network size and social structure category. (a,b) Density decreased with network size and was reduced in the multi-level societies, while (c,d) modularity increased with network size and was elevated in both the cliquish and multi-level societies. Points indicate mean values for each species, while error bars indicate uncertainty (± 2 SE) in both network size and global network metrics, pooled across the 20 posterior networks for each group-year. Shaded intervals indicate the 89% credible intervals from the zero-one-inflated beta regression models examining the impacts of network size and social category on network metrics. Points and ribbons are colored according to their respective social category. Shaded intervals are truncated along the x-axis to the minimum and maximum network sizes observed within each respective category.

In general, grooming network metrics from distinct populations of the same species were generally similar, suggesting that network structure is a species-level trait. However, because only two species were represented by multiple populations, our dataset likely underestimates intraspecific variation. Also, network metrics did not map neatly onto the phylogeny of the papionins (**Figure 1b**). Intraclass correlation coefficients (ICC) from network size-only models indicate that only 15-21% of the variance in these global network metrics could be attributed to phylogenetic position (Density: ICC_Phylogeny_ = 0.154, ICC_Species_ = 0.138, ICC_Population_ = 0.052; Modularity: ICC_Phylogeny_ = 0.206, ICC_Species_ = 0.237, ICC_Population_ = 0.057). In other words, while closely related species sometimes formed similar grooming networks, more distantly related species occasionally did so as well. We explore the social forces (e.g., the strength kin and rank biases, the extent of male clustering effects) that could generate these convergent similarities in the analyses below.

### Female-female grooming ties were variably shaped by kinship, rank, and shared male ties

To identify the patterns that produce variation in grooming network structure, we constructed a set of models focused on three factors that often shape female-female relationships: (i) maternal kinship (known, estimated, or imputed *r*_*mat*_ *≥* 0.25; yes/no); (ii) rank similarity (i.e., 1 - the absolute difference in their proportional ranks); and (iii) shared male ties, defined as whether the two females shared the same top male grooming partner (given plus received) during that group-year. For most populations, maternal kinship was determined from known maternities. By contrast, because data from Guinea baboons depended on genetic sampling alone, we could not exclude paternal kin from these data, and caution is warranted when comparing this population with the others. When we were unable to determine maternal relatedness, kinship was imputed using informative criteria (13.3% of dyads, see *Supporting information* for quality control tests). Based on previous work, we expected females to selectively groom their relatives and other closely ranked groupmates (38–40, 44). However, we expected kin and rank biases to be diluted in species with female-biased dispersal (45, 46). In the multi-level societies, female-female relationships may largely be a byproduct of shared male ties, as females primarily socialize within their one-male units (54, 78). However, similar if weaker shared male effects could be present in the single-level papionins, in which shared male ties have been linked to strengthened female-female proximity relationships (e.g., olive baboons: 51). If so, then these propensities could be the evolutionary precursors of multi-level units.

To test these predictions, we constructed regression models examining the impact of these three factors on undirected grooming patterns within each group-year (n = 126 annual networks, excluding group-years with <4 females or <2 males), using 20 draws from the posterior distribution of edge weights to propagate uncertainty in grooming relationships. To normalize edge weights, we divided each undirected edge weight by the mean value observed for that group-year and analyzed these standardized relationship strengths using a ‘Gamma’ distribution. However, when measuring dyadic effects in hamadryas baboons, we took an alternative approach. First, we excluded kinship as a covariate, as relatedness is often obscured by female transfer and genetic sampling postdates the behavioral sample (45). Second, we relied on randomized female rank values, as contests between female hamadryas are mostly undecided (54) and thus rank effects are expected to be negligible.

We then extracted effect sizes and standard errors from these models and constructed a series of Bayesian phylogenetic meta-analyses examining whether the magnitude of these effects varied as a function of social category. To account for the impacts of demography on these interaction patterns, we also included network size and female dispersal regime (i.e., female philopatry vs. female-biased dispersal) as moderators. To account for repeated measures, these models incorporated random effects for phylogeny, species, population, and group identity.

Kinship was a robust predictor of female-female grooming relationship strengths across all three social categories (**Figure 3a**). Within the single-level societies, kin biases were salient in the cohesive societies (*θ*_Cohesive_ = 1.29, 89% CI = [0.92, 1.63]) but slightly stronger in the cliquish societies (*θ*_Cliquish_ = 1.72, 89% CI = [1.25, 2.14]). Within the multi-level societies, kinship effects were stronger in female-philopatric geladas than in female-dispersing Guinea baboons (β_Dispersal_ = -1.08, 89% CI = [-1.99, -0.13]). Kin biases also tended to be stronger in larger grooming networks (β_GroupSize_ = 0.51, 89% CI = [0.34, 0.67]). Rank similarity effects were weak in the cohesive societies (*θ*_Cohesive_ = 0.17, 89% CI = [0.04, 0.30]) but moderate in the cliquish societies (*θ*_Cliquish_ = 0.49, 89% CI = [0.33, 0.64]). By contrast, rank effects were undetectable in the multi-level societies in which rank information was available (*θ*_Multi-level_ = -0.05, 89% CI = [-0.30, 0.20], **Figure 3b**). While sharing the same top male did not predict the strength of female-female grooming relationships in the cohesive and cliquish societies, this effect was pronounced in the multi-level societies (*θ*_Multi-level_ = 2.45, 89% CI = [1.93, 2.93], **Figure 3c**), in which females residing in the same reproductive units often shared the same top males by default.

**Figure 3.**
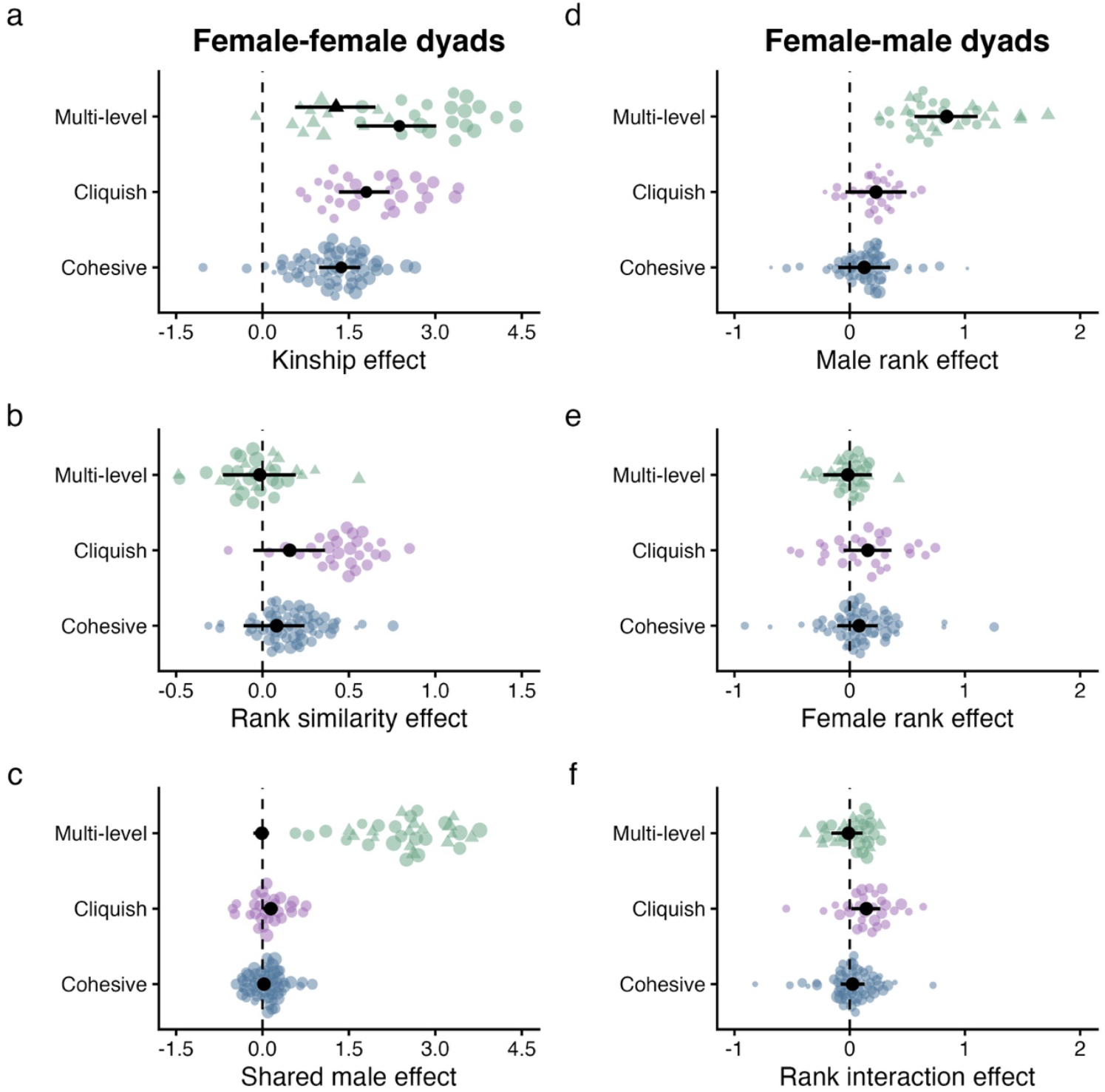
The strength of female-female nepotistic biases and female-male rank biases varied across species. In (a,b,c) circles indicate species with female philopatry, and triangles indicate species with female-biased dispersal. (a). Kinship was a salient but variable predictor of female-female grooming relationships. Here, we provide a separate posterior estimate for species with female-biased dispersal, as this was the only case where dispersal patterns had a meaningful impact on the effect size of interest. (b) Rank similarity effects were present in cohesive societies, strongest in cliquish societies and undetectable in multi-level societies. (c) Females that shared the same top male partners formed stronger grooming relationships in the multi-level groups but not in the single-level groups. (d) High-ranking and reproductively dominant males received more grooming from females, and this tendency was far more salient in the multi-level societies. (e) High-ranking females did not typically form stronger between-sex relationships. (f) Rank interaction effects were likewise weak-to-absent across all three groups. Each point indicates a standardized effect size from one group-year within the database. In (b), (d), and (e), hamadryas baboons were excluded for visualization purposes, as female contests were undecided and therefore ranks randomized. However, models including and excluding these presumed near-zero effects were broadly equivalent. Black points and error bars indicate posterior means and 89% credible intervals. The size of each point is scaled to the effect’s precision (1/SE).

### Female-male grooming ties were variably shaped by social dominance

We then examined the effects of dominance rank on the strength of female-male grooming relationships, as female and male ranks can both predict between-sex grooming patterns (48, 50, 79). Specifically, we aimed to quantify undirected female-male grooming interactions as a function of: (i) male rank; (ii) female rank; and (iii) their interaction. Including an interaction term allowed us to capture whether high-ranking males have priority of access to high-ranking females and vice versa (i.e., rank assortment), which could reflect bidirectional competition for mating partners. Because males within multi-level groups do not show conventional dominance relationships (64, 80), we labeled leader males as high-ranking (rank = 1) and follower males as low-ranking (rank = 0). While these rank metrics correspond with distinct social phenomena across the single-level and multi-level groups, they both reflect patterns of mating behavior and paternity success (40, 81–86). As above, we calculated standardized edge weights and constructed regressions similar to those above for each group-year in the data set (n = 124, excluding group-years with <3 males). Then, we extracted effect sizes for the predictor variables and conducted phylogenetic meta-analyses to determine whether the strength of these effects varied as a function of social category and network size.

Across the sample, females tended to groom with high-ranking males more than with low-ranking males. However, these patterns were not universal. Male rank effect credible intervals overlapped zero across the single-level societies (*θ*_Cohesive_ = 0.13, 89% CI = [-0.10, 0.35]; *θ*_Cliquish_ = 0.23, 89% CI = [-0.04, 0.50]) but were far from zero in the multi-level societies (*θ*_Multi-level_ = 0.84, 89% CI = [0.56, 1.11], **Figure 3d**). Thus, male rank effects are weaker and more inconsistent when males form linear hierarchies but stronger and more regular when dominant males wield substantial social and reproductive leverage. Across the papionins, female dominance rank had no consistent relationship with the strength of female-male grooming relationships (**Figure 3e**). Rank interaction effects often emerged in the cliquish papionins, although credible intervals indicate relatively weak support for this rank assortment effect (*θ*_Cliquish_ = 0.14, 89% CI = [0.01, 0.26]; **Figure 3f**).

## DISCUSSION

Using a rich, new comparative dataset, we captured wide variation in group structure and its underpinnings across the papionins. As expected, species that live in single-level societies formed groups that were denser and less modular than species that live in multi-level societies. However, consistent with qualitative distinctions made in recent work (49), the single-level species could be divided into two distinct categories. Mandrills, Kinda baboons, and chacma baboons (‘cliquish’ societies) formed grooming networks that were consistently more modular than those of yellow baboons, olive baboons, Sanje mangabeys, sooty mangabeys, and gray-cheeked mangabeys (‘cohesive’ societies). In turn, these three social categories mapped onto variation in the social factors that shaped the strength of female-female grooming relationships. While kin and rank biases were salient across the single-level societies, these tendencies were elevated in cliquish groups. Shared male ties played a central role only in the multi-level species, suggesting that this pattern may be exclusive to these social systems. Likewise, while female-male grooming relationships were often strong, male rank effects were only prevalent in multi-level groups. Below, we discuss how these social strategies may be tailored to distinct social and ecological pressures and propose comparative tests that might uncover the proximate and ultimate drivers of this variation.

### Why do the single-level societies vary in their structure?

While group size was generally higher in the cliquish species than in the cohesive species, differences in their grooming network measures persisted even after accounting for network size. These differences could, at least in part, be a byproduct of papionins’ evolutionary history. Cliquish chacma baboons and Kinda baboons are closely related and share a history of extensive admixture (70, 87–89). However, phylogenetic similarity cannot explain the evolution of cliquish grooming networks in distantly related mandrills, suggesting an additional role of social or ecological pressures in generating convergence across disparate lineages. Harsh and heterogeneous environments could induce increased female-female competition over food resources and encourage females to prune their social ties (90, 91) and form cliquish subgroups (59). Forthcoming analyses using the CAPS database will test this ecological hypothesis by linking variation in network structure with key measures of environmental productivity and habitat heterogeneity. However, this is unlikely to be the primary explanation, as cliquish species do not universally occupy harsh or seasonal environments (92, 93), and papionin groups do not always become more fragmented during resource-poor conditions (94–96).

Instead, the heightened modularity we observed in cliquish groups could reflect competition over male partners. Males within the cliquish societies attracted more grooming than males in the cohesive societies (**Figure S1**), and all three cliquish taxa show marked – if contrasting – signatures of sexual selection. Both chacma baboons and mandrills exhibit high canine and body size dimorphism (97), and male mandrills further display conspicuous facial coloration (98). These characteristics correspond with heightened contest competition over mating opportunities (99, 100), high levels of male reproductive skew (40, 82, 101), and, perhaps, elevated infanticide risk (93, 102), which could incentivize females to form relationships with likely sire males in order to protect their vulnerable offspring (50, 103). By contrast, Kinda baboons have high relative testes sizes and exhibit mating patterns that are indicative of indirect male-male competition (104, 105), suggesting that strong female-male relationships in this species may be the result of mate choice and may provide different adaptive benefits (e.g., improved offspring development: 106). In both of these systems, limited male attention could exacerbate female-female competition, thereby heightening female social selectivity. Alternatively, strong between-sex bonds could generate time budget constraints, preventing females from grooming all of their female groupmates. If so, then apparent social selectivity could emerge rather as a byproduct of these altered social priorities. Future analyses using the CAPS datasets will thus target the drivers of strong, stable between-sex bonds (reviewed in 15) and model their wider impacts on network structure.

### Kinship and rank variably predicted the strength of grooming relationships

As expected, maternal kinship was a robust but variable predictor of grooming ties across papionin taxa. Much of the variation in kin bias was attributable to network size and female dispersal patterns, likely because variance in relatedness increases when groups become larger and habitual female dispersal typically reduces the availability of kin (107, 108). However, variation in relatedness patterns does not always explain differences in social structure across groups (e.g., hyenas: 109, macaques: 110) and between related species (e.g., bats: 111). Indeed, categorical differences in kin bias persisted even after accounting for these known drivers of local relatedness. Specifically, kin biases were strongest in multi-level geladas, strong in the cliquish societies, moderate in the cohesive societies, and moderate-but-variable in the multi-level Guinea baboons, in which females disperse from their natal units (46). Thus, all else being equal, the stronger kinship effects observed in the cliquish papionins may reflect a more selective social environment (e.g., investment in relationship quality over quantity: 112).

Dominance rank effects were similarly widespread but highly variable across the papionins. In general, female-female dyads that were close in rank tended to groom more than dyads that were further apart in rank. Although kin also tend to be closer in rank (due to maternal rank inheritance: 43, 113, 114), these rank similarity effects persisted even after accounting for kinship (as in 38, 39, 44). In general, rank similarity effects were weak in the cohesive societies but stronger in the cliquish societies, which may be marked by more intense forms of female-female competition. Specifically, data on aggression patterns suggest that cohesive olive baboon females compete primarily over food resources (57, 115), while cliquish chacma baboon females compete more intensively over male partners (50, 103, 116). If male attention is more limited and less easily shareable than food, this difference could help explain the stronger rank effects observed in cliquish systems. Rank effects were generally absent in the multi-level societies, perhaps because rank is more salient within one-male units (e.g., 44) than in the higher social levels captured in our analyses. However, competition appears to be broadly blunted in multi-level groups (54, 94, 95, 117, 118), so these patterns may reflect a reduced importance of social hierarchies in multi-level societies.

Rank effects on female-male relationships were also present but surprisingly inconsistent. While high-ranking males tended to receive more grooming than low-ranking males, this pattern only received strong empirical support in the multi-level societies. Thus, male rank effects might only become salient when males wield substantial social and reproductive control (15, 56). In multi-level societies, leader males hold near-exclusive access to the females within their units (52, 53, 119). By contrast, in single-level societies, low- and mid-ranking males can more reliably secure reproductive opportunities (120), often by forming strong bonds with particular females (83). Female rank had no association with female-male grooming, suggesting that papionin females do not uniformly compete for access to males. However, rank assortment patterns (e.g., 79, 121) were weakly present in the cliquish papionins. Thus, female-female competition over high-quality mates may emerge in some circumstances, particularly when males are poised to provide anti-infanticide protection or other offspring benefits (50, 103). In combination, these data suggest that dominance rank effects are highly variable across species, likely because females compete over different resources (e.g., access to food vs. protective males: 115, 118, 122) and males face varying degrees of mate competition (as indexed by reproductive skew: 40, 81–85).

### Shared male effects were limited to the multi-level societies

Females in multi-level societies primarily groomed females from their own reproductive units (44, 52, 54), generating strong shared male effects. These patterns were similarly pronounced across all three multi-level taxa, despite variation in the observed degree of female social agency: female Guinea baboons move freely between units (52), female geladas typically remain with their valuable kin partners (43), and female hamadryas are largely restricted by their leader males (65). Although baboon females who share the same preferred male occasionally form stronger spatial associations (51), shared male effects were negligible across the single-level papionins. In other words, shared male preferences might bring females into close proximity without prompting the formation of strong grooming relationships. In this way, the between-sex bonds documented in the single-level societies (48, 50, 103, 123) are largely distinct from those found in the multi-level societies (52–54), as the latter are more integral to grooming network structure (124).

More broadly, these patterns suggest that the social forces at play in the semi-modular cliquish societies (i.e., kin and rank biases) are largely distinct from those present in the highly modular multi-level societies (i.e., shared male effects). That being said, only two species were represented by multiple study populations. Thus, the CAPS sample likely underestimates intraspecific variation in network structure and may not capture all the ways in which single-level societies overlap the multi-level societies. For instance, none of the cliquish chacma baboon groups sampled here formed one-male-unit-like subgroups (59), and the sampled Kinda baboon group was much smaller than the average identified in wider population censuses (104). Including more study populations could alter our evolutionary inferences. Specifically, because chacma and Kinda baboons have both been positioned as basal members of *Papio* in genomic analyses (87, 125), increased sampling of these species could uncover ancestral social phenotypes that might have facilitated the later evolution of modular social structures in the hamadryas and Guinea baboon lineage but were seemingly lost in yellow and olive baboons.

The ancestral condition for geladas, however, is perhaps more ambiguous. While ancestral *Theropithecus* may have formed large, baboon-like groups, small and cohesive groups (like those of *Lophocebus*, gray-cheeked mangabeys) cannot be ruled out using phylogenetic principles (29). If so, then small, ancestral *Theropithecus* groups could have coalesced into the large supergroups observed in extant geladas (more closely mirroring the evolutionary pattern inferred for multi-level Asian colobines: 126). In future work, it may be useful to shift the scale of network analysis (i.e., the unit or the band) when quantifying gelada social structures, thereby identifying which level of their social system is most likely homologous to the single-level group. Regardless, the differing patterns we documented here underscore the fact that – despite their structural commonalities – multi-level societies are diverse in their overall shape, presumed functions, and evolutionary histories (28, 127).

## CONCLUSIONS

This new dataset provides a systematic foundation for mapping behavioral variation across the papionins and identifying the forces that produce between-species differences in social structure. Taken together, these results demonstrate that the papionins are more socially diverse than has been historically recognized. Beyond the well-known distinction between species that form single-level and multi-level societies, our analyses show that single-level societies can be further divided into two social categories, cohesive and cliquish. Kin biases widely influenced females’ grooming behavior across the papionins, but female-female rank biases were markedly elevated in the cliquish species. Additionally, shared male patterns were only pronounced in the multi-level societies, suggesting that these propensities may be exclusive to these species. It remains unclear, however, why natural selection has favored these particular strategies or, more broadly, why multi-level societies have evolved in several taxa, including humans (28, 127). Further analyses using the CAPS database – alongside ongoing genomic studies (87, 125, 128–130) and a growing papionin fossil record (131, 132) – may provide novel, complementary insights into the evolution of modular social structures in papionins and other radiations.

## MATERIALS AND METHODS

### Study sites and populations

The data used for this study were collected across multiple wild populations of African papionins (see **Table S1** for a list of projects contributing to CAPS). Details regarding the demographic histories and data collection methods across these long-term research projects can be found in previous reports (40, 41, 44, 48, 65, 86, 104, 122, 133–138). Each contributing study provided the records necessary to construct most or all of the following data frames: (i) study subjects; (ii) group membership; (iii) dates and durations of behavioral observations; (iv) grooming interactions; (v) dyadic kinship; (vi) female and male ranks, aggregated as yearly mean values for each individual. Below, we describe how these datasets were generated, how this information varied across sites, and how we treated missing information.

### Study subjects and group membership

Contributing projects provided either (i) known and/or estimated dates of birth or (ii) age categories (e.g., subadult, young adult, etc.) for all adult female and male subjects that were included in behavioral sampling. In total, the sample comprised 1207 unique adults and subadults (727 females, 480 males) residing within 28 distinct social groups (1-5 groups per population, **Table S1**). When constructing grooming networks, we included females if their known or estimated age was ≥ 4 years (when females begin to mature: 139, 140–142) or if they were classified as subadults or adults during that calendar year. Likewise, we included males if their known or estimated age was ≥ 7 years (when males reach life history milestones: 141, 143–145) or if they were classified as subadults or adults during that year. To determine periods of co-residence among each pair of individuals, we used group membership data from each study site, noting when individuals ‘entered’ (due to their birth or immigration) or ‘exited’ (due to their death, disappearance, or dispersal) their respective social groups. This allowed us to exclude behavioral samples collected when pairs were no longer co-resident when calculating dyadic observation effort. Most individuals belonged to one social group throughout the study periods. However, males (n=47) occasionally dispersed from one study group into another, and females (n=10) occasionally switched social groups via dispersal (in Guinea baboons) or during group fission events (in the olive baboons at Laikipia).

### Behavioral observations and grooming interactions

Each contributing study site provided behavioral observation records at the individual-level. In most cases, behavioral data were collected via continuous focal animal observations (5-675 minutes in duration, mean = 19.8 minutes), during which the direction and duration of the focal animal’s grooming interactions were recorded. However, at some study sites (Amboseli and Gashaka, **Table S1**), durationless grooming events were collected opportunistically while researchers moved about the study group and collected other routine data (e.g., focal samples). This method of data collection, termed representative interaction sampling, and associated procedures used to correct for sampling densities at the group-level have been described elsewhere (146, 147). In short, because grooming records extended beyond the focal individual, observation effort was calculated not at the individual-level but at the group-level (i.e., mean number of focal samples per female). For one population (hamadryas baboons at Filoha), social interactions were recorded during unit-level scans collected at ∼10-minute intervals (54).

### Kinship estimates

Kinship was determined using a mixture of maternal pedigree information (available from all sites to varying depths), genetically confirmed maternities (Simiens: 148, Amboseli: 149), and genetic relatedness estimates (Simenti: 150). Using this information, we estimated maternal relatedness for all dyadic pairs and designated dyads as being related if *r*_*mat*_ ≥ 0.25 (e.g., mother-offspring and maternal sister dyads). However, because kin relationships among Guinea baboons at Simenti are only known from genetic sampling, dyads with *r* ≥ 0.25 may include paternal kin. Thus, kin bias estimates in this species could differ solely due to methodological reasons. Given differences in pedigree depth and genetic sampling, sites varied widely in the relative representation of related, unrelated, and unknown dyad pairs. To avoid potential biases and to improve inferences for studies with shallower pedigrees, we implemented classification and regression tree imputation to categorize unknown dyads using informative criteria (such as the dyad’s rank similarity, see *Supporting information*). However, we did not impute kinship data for hamadryas baboons and excluded kinship from analyses focused on this population, as both genetic and observational data were limited. When conducting analyses focused on female-male dyads, we removed related pairs, as kin dyads are unlikely to form mating relationships (e.g., 151).

### Dominance ranks

Dominance ranks were (i) directly provided by the collaborating research projects or (ii) calculated from directed agonistic interactions using the ‘EloRating’ package (152). To standardize dominance ranks within each study group, we calculated rank as the proportion of all same-sex group members that each individual outranked. However, given differences in the scale of competition across the multi-level societies, we calculated ranks differently in these three species. Specifically, female ranks in geladas were calculated at the level of the reproductive unit (43), as between-unit aggression is rare and competition is highly localized, while female ranks in Guinea baboons were calculated at the party-level (117). By contrast, contests between female hamadryas baboons were generally undecided, so dominance ranks were inestimable (54). Thus, we imputed random dominance ranks for each female and recovered a presumed null effect. Likewise, males in multi-level societies do not form clear dominance hierarchies across their entire ‘bands’ or ‘parties.’ Thus, we categorized leader males as having ranks of 1, and follower males as having ranks of 0.

### Grooming network construction

Annual group-level grooming networks were constructed in ‘bisonR’ (66) and visualized using ‘igraph’ (153). Edges were calculated using (i) the total amount of time that a dyad engaged in grooming (in minutes, rounded upwards), modeled using a binary conjugate prior; or (ii) the number of grooming events observed during ‘representative sampling’ (147), modeled with a count conjugate prior. To account for differences in sampling effort, we treated the total duration of focal observation or the number of instantaneous scans as an ‘exposure term.’ This allowed us to account for differences in observation effort and propagate edge weight uncertainty. After removing edge weights that fell below the minimum observed value, we then calculated two metrics from each posterior network: (i) density, which captures the proportion of dyads that were observed grooming; and (ii) modularity, which captures the tendency for individuals to groom within clusters or subgraphs that are determined using a random walk algorithm (‘cluster_walktrap’).

### Statistical analyses

First, to quantify the drivers of grooming network density and modularity, we constructed two models using the ‘brm_multiple’ function in ‘brms’ (71), pooling effects across 20 posterior networks from the ‘bisonR’ edge models. We modeled density and modularity using a zero-one-inflated beta distribution, as these network values are bounded between 0 and 1, and included network size and social category (i.e., cohesive, cliquish, and multi-level) as fixed effects. To account for phylogenetic history, we included a random effect using a genomically informed phylogenetic covariance matrix (70). Because the three mangabey taxa were not included in this phylogeny, we used closely related congeners that match their relative phylogenetic position. We also included species, population, and social group identities as nested random effects. This approach allowed us to attribute variation in network structure to (i) phylogenetic history, (ii) species-specific traits that are independent of their phylogenetic history, and (iii) population- and group-level effects.

Second, to determine the factors that underpin social relationships across populations, we constructed a series of dyadic regression models. For each group-year in the dataset (n = 126, excluding group-years with <4 females or <2 males), we constructed a regression in which standardized, undirected edge weights were treated as ‘Gamma’ outcomes, as posterior predictive checks indicated that this family adequately captured the skew in these data. In each model, we included kinship (yes/no), rank similarity (i.e., 1 - the dyad’s absolute proportional rank difference), and shared top male (yes/no) as fixed effects. When rank data were missing (because a female had yet to enter the dominance hierarchy or her Elo-rating had yet to stabilize, etc.), we included her rank from the first year she was included in the dataset. For hamadryas baboons, we assigned females randomized ranks, as contests are undecided in the wild (54). Removing effect sizes that relied on these randomized ranks did not qualitatively change our results. We included actor and recipient identities as multi-membership random effects to account for nodal interdependence. As noted above, covariates were missing for some dyads. Thus, we used classification and regression tree imputation using the ‘mice’ package (154) to impute missing individual attributes (proportional ranks) and dyadic covariates (kinship) based on informative criteria. Critically, imputing these missing data did not bias our effect size estimates (see *Supporting information*). For each group-year, we pooled model estimates across 20 imputed datasets using the ‘brm_multiple’ function in ‘brms’ (71)

We used a similar process when modeling the predictors of female-male relationship strengths, using female proportional rank, male proportional rank, and their interaction as fixed effects. We again included randomized ranks for hamadryas baboon females. As above, we used imputation procedures to include males with missing rank information and pooled model estimates across 20 imputed datasets for each group-year. For each group-year in the dataset (n = 124, excluding group-years with <3 adult or subadult males), we constructed a ‘Gamma’ model similar to the one outlined above, including actor and recipient as multi-membership random effects. We then extracted effect size estimates and standard errors for use in downstream meta-analyses.

To assess differences in these social processes across the sample, we then constructed six phylogenetic meta-regressions in ‘brms’ to assess the dyadic effects listed above (female-female dyads: kinship, rank similarity, and shared male effects; female-male dyads: female rank, male rank, and their interaction). Each effect size was treated as a student-distributed outcome, as this better accommodated highly skewed effect sizes distributions. We incorporated standard errors to account for measurement uncertainty and included social category (cohesive, cliquish, multi-level), network size, and female philopatry (yes/no, female-female effects only) as fixed effects in each of these models. We also controlled for phylogeny, species, population, group identity, and group-year using random effects. To verify the robustness of these results (155), we performed frequentist meta-analyses using ‘metafor’ (156). The output of these models was largely concordant with those using the Bayesian approach. Across all models, we standardized continuous variables for improved interpretation. Rhat values were low across all models (≤1.01), and posterior predictive checks indicated adequate fits for network metrics, dyadic data, and meta-regression estimates. To avoid conflation between our Bayesian model outputs and *P-*values, we provide 89% credible intervals when reporting and visualizing uncertainty in our estimates (157).

## DATA AVAILABILITY

The data and code necessary to reproduce these analyses and associated visualizations are available on GitHub (https://github.com/PapioninSocieties/Feder_et_al_CAPS_2025) and have been archived on Zenodo (https://doi.org/10.5281/zenodo.16422722).

## ACKNOWLEDGEMENTS

We thank the many researchers and field assistants who have collected, collated, and managed behavioral and demographic data across the projects that contributed data to the Comparative Analysis of Papionin Societies (CAPS). We would also like to thank the many funding agencies and wildlife research organizations that have supported long-term research at these study sites. Specific acknowledgements for each contributing research project can be found in the *Supporting information*. The creation of the CAPS database and the writing of this manuscript were supported by an NSF SBE Postdoctoral Research Fellowship (SMA-2313739, to JAF).

