## Supporting information for "Disparate social structures are underpinned by distinct social rules across a primate radiation"

### Comparative Analysis of Papionin Societies
### Contributing Study Acknowledgements

**Amboseli Baboon Project**
We gratefully acknowledge the support of the National Science Foundation and the National Institutes of Health for the majority of the data represented here, most recently through R01AG071684, R01AG075914, and R61AG078470. Current support for field-based data collection also comes from the Max Planck Institute for Evolutionary Anthropology, and we thank Duke University, Princeton University, and the University of Notre Dame for financial and logistical support. In Kenya, our research was approved by the Wildlife Research Training Institute (WRTI), Kenya Wildlife Service (KWS), the National Commission for Science, Technology, and Innovation (NACOSTI), and the National Environment Management Authority (NEMA). We also thank the University of Nairobi, the Institute of Primate Research (IPR), the National Museums of Kenya, the members of the Amboseli-Longido pastoralist communities, the Enduimet Wildlife Management Area, Ker & Downey Safaris, Air Kenya, and Safarilink for their cooperation and assistance in the field. Particular thanks go to the Amboseli Baboon Project long-term field team (R.S. Mututua, S. Sayialel, J.K. Warutere, I.L. Siodi, I.L., and L. Musembei), and to T. Wango and V. Oudu for their untiring assistance in Nairobi. The baboon project database, Babase, is expertly managed by J. Gordon, and W. Wilbur, with past assistance by N. Learn. Database design and programming are provided by K. Pinc. This research was approved by the IACUC at Duke University, University of Notre Dame, and the Ethics Council of the Max Planck Society and adhered to all the laws and guidelines of Kenya. For a complete set of acknowledgments of funding sources, logistical assistance, and data collection and management, please visit http://amboselibaboons.nd.edu/acknowledgements/.

**CRP Simenti**
We want to thank the Direction des Parcs Nationaux (DNP) and the Ministère de l'Environnement et de la Protection de la Nature (MEPN) du Sénégal for approval to conduct this study in the Parc National du Niokolo-Koba (PNNK). This research was funded by the Deutsche Forschungsgemeinschaft (DFG, German Research Foundation) - Project-ID 254142454/GRK 2070 "Understanding Social Relationships".

**Filoha Hamadryas Project**
The Filoha Hamadryas Project would not be possible without permission and facilitation from the Ethiopian Wildlife Conservation Authority (EWCA), and in this respect we are especially grateful to Kumara Wakjira, Kahsaye Gebretensaye, Girma Ayalew, and Fanuel Kebbede. Thanks also to the scouts and wardens of Awash National Park and the students, collaborators, field managers, and field assistants who have worked with the project over the past three decades. Fieldwork at Filoha during the time these data were collected was funded by the Leakey Foundation (1998), the Wenner-Gren Foundation (1996), the National Geographic Society (6468-99), and the National Science Foundation (9629658).

**Gashaka Baboon Project**

This work would not have been possible without the initial baboon habitation and data collection by Ymke Warren (1970-2010) and Jeremiah Adanu (1969-2004), and the tireless efforts of Volker Sommer who set-up and led the Gashaka Primate Project. Many students and volunteers contributed to the data set used here- including Nienke Alberts, David Inglis, Gonçalo Jesus, Emily Lodge, and a number of MRes and MSC students from Roehampton University and University College London. Ann MacLarnon and Stuart Semple provided student supervision and project input over many years. We are very grateful to field assistants Bobbo Buba, Halidu Iliyasu, Hammaunde Guruza, Ibrahim Usman, Maikanti Hassan, Maigari Ahmadu, Buba Hammaselbe and Felix Vitalis, who were indispensable in the collection of field data. The Gashaka Baboon Project was supported by funding from Roehampton University. Fieldwork was enabled by permits from the Nigeria National Parks Service to the Gashaka Biodiversity Project (previously Gashaka Primate Project), and core funding from the North of England Zoological Society/ Chester Zoo Nigeria Biodiversity Program.

**Kasanka Baboon Project**

We thank the Zambian Department of National Parks and Wildlife (DNPW) and Kasanka National Park for research collaboration and permission to conduct fieldwork in Kasanka National Park. We thank Marley Katinta and Kennedy Kaheha for their long-term work as wildlife scouts and researchers. We thank the numerous camp managers for their support of data collection including Elizabeth Winterton, Aileen Sweeney, Cassandra Ekhdahl, Kim Gordon, Rachel Sassson, and Roxanne de Rodez-Bénavent. Thank you to Frank Willems and Inge Akerbom for initial and continued field project support and logistics. Data used in this manuscript was supported by the Fulbright Program, Washington University in St. Louis, Lambda Alpha, American Society of Primatologists, P.E.O. International, Idea Wild, the John W. and Helen B. Jarman Foundation and the University of Massachusetts.

**Kibale Mangabey Project**

We thank the Uganda Wildlife Authority, Uganda National Council for Science and Technology, and personnel at the Makerere University Biological Field Station in Kanyawara for permission to work in Kibale National Park. We thank all the field assistants that worked with us during these years: Kaseregenyu Richard, Katusabe Swaibu, Irumba Peter, Sabiti Richard, Akora Charles, and Koojo John for their invaluable assistance in the field. This research was supported by funds from the NIH/NIA grants PO1 A6022500 and PO1 A608761 (to J.R. Carey) and “Mobilitas” postdoctoral grant MJD56. This study complies with the current laws of Uganda.

**Laikipia Baboon Project**

We thank the Office of the President of the Republic of Kenya and the Kenya Wildlife Service for permission to conduct this field research. We thank Dr. Shirley Strum for her permission to work with the Uaso Ngiro Baboon Project and for access to long-term data. We thank Kate Aberholden, Megan Best, Megan Cole, Moira Donovan, Alexandra Duchesneau, Jessican Gunson, Molly McEntee, Laura Pena, and Leah Worthington for their contributions to data collection, Eila Roberts for supervising the field project, and the staff of the Uaso Ngiro Baboon Project, particularly Jeremiah Lendira, Jane King’au, Joshua Lendira, and Frances Molo, for providing support in the field. We also thank David Muriuri, who provided invaluable assistance with logistics and data management, and the African Conservation Center, which facilitated the project. The field project was supported with funding from Arizona State University to J.B.S.

**Lékédi Mandrillus Project**
The Mandrillus Project is extremely grateful to all the past and present field assistants for their daily data collection since 2012. We also thank the SODEPAL-COMILOG society (ERAMET group) for their long-term logistical support. The Mandrillus Project has been funded by several grants that allowed long-term data collection including: SEEG Lekedi, MITI and SEE-LIFE initiative (INEE-CNRS), the Leakey Foundation (S202210309), and the Max Planck Society (all to MJEC). Data collection was approved by an authorization from the CENAREST institute (permit no. AR15/24/MESRSIT/CENAREST/CG/CST/CSAR). This is a Project Mandrillus publication number XXX and ISEM-SUD 2025-XXX.

**Moremi Baboon Project**
We would like to thank the Office of the President of the Republic of Botswana and the Botswana Department of Wildlife and National Parks for permission to conduct research on chacma baboons in the Moremi Reserve. We also thank Dorothy Cheney and Robert Seyfarth for providing access to long-term life history and behavioral data. Data on chacma baboons were collected by Jacinta Beehner, Thore Bergman, Dorothy Cheney, Catherine Crockford, Anne Engh, Julia Fischer, Marlies Heesen, Dawn Kitchen, Liza Moscovice, Ryne Palombit, Drew Rendall, Keena Seyfarth, Robert Seyfarth, Chantelle Shaw, Joan Silk, Roman Wittig, Mokopi Mokopi, and Alec Mokopi. Research in Moremi was supported by grants from the National Science Foundation (IOS-9514001), the National Institutes of Health, the DFG and KFN, the NSERC of Canada, the National Geographic Society, the Leakey Foundation, and the University of Pennsylvania.

**Simien Mountains Gelada Research Project**
We would like to thank the Ethiopian Wildlife Conservation Authority for permission to conduct research on geladas in the Simien Mountains National Park (SMNP), Ethiopia. We would also like to thank the wardens of the SMNP and the many field team members that collected these data, in particular Eshete Jejaw, Ambaye Fenta, Setey Girmay, Julie Jarvey, Megan Gomery, Levi Morris, Tara Regan, Patsy DeLacey, Peter Clark, Evan Sloan, Liz Babbitt, and Maddie Melton. This work was supported by the National Science Foundation (BCS-0715179 to TJB and JCB, BCS-1723228 to AL, IOS-1255974 to JCB and TJB, IOS-1854359 to JCB and TJB, BCS-1723237 to NSM, BCS-2010309 to NSM, BCS-2017976 to JAF), the Leakey Foundation, the National Geographic Society (NGS-8100-06, NGS-8989-11, NGS-1242, and NGS-50409R-18), the Fulbright Scholars Program, the University of Michigan, Stony Brook University, and Arizona State University.

**Taï Chimpanzee and Mangabey Project**
We thank the Ministère des Eaux et Forêts and the Office Ivoirien des Parcs et Réserves in Ivory Coast for permitting this data collection (Research permit: Wittig/006/MESRS/DRI). Special thanks to Christophe Boesch for establishing and nurturing the Taï Chimpanzee Project, the Centre Suisse de Recherches Scientifiques, field assistants, and all members of the Taï Chimpanzee and Mangabey Project for their invaluable support.

**Tsaobis Baboon Project**
We are very grateful to the Tsaobis Baboon Project field assistants from the 2005–2006 and 2013–2014 seasons for their invaluable contributions to data collection. We thank the Swart family, the Ministry of Lands and Resettlement of Namibia, and the Tsaobis beneficiaries for permission to work at Tsaobis; the Snyman and Wittreich families, Desert Craft, and J. Venter for permission to work on

their land; and the Gobabeb Namib Research Institute for affiliation. This research was carried out with the authorisation of the Ministry of Environment and Tourism of Namibia (MET Research/Collecting Permits 886/2005, 1039/2006, 1786/2013, 1892/2014). E.H. and A.B. benefited from financial support from the French ‘Ministère de l'Enseignement Supérieur, de la Recherche et de l'Innovation’. G.C. was supported by the UK’s Natural Environment Research Council and Research England. This paper is a publication of the ZSL Institute of Zoology’s Tsaobis Baboon Project. Contribution ISEM-SUD 2025-XXX.

**Udzungwa Sanje Mangabey Project**
Permission to carry out research on Sanje mangabey was granted by the Tanzania Commission for Research and Technology, Tanzania National Parks, and Tanzania Wildlife Research Institute. We thank the Udzungwa Mountains National Park staff and the Udzungwa Ecological Monitoring Centre staff, who provided invaluable logistical support. We also thank Thad Bartlett, Carola Borries, Janine Brown, Diane Doran-Sheehy, John Fleagle, Jessica Rothman, Patricia Wright, and especially Carolyn Ehardt, who set up the Sanje Mangabey Project. Special thanks to the Sanje Mangabey Project research team: Saidi Amili, Ally Chitita, the late Amani Kitegile, Amos Lumagi, Clever Ngatwika, Fracis Masinde, Salimini Saidi, Baraka Sehaba, Aloice Mwakisoma and especially Yahaya Sama, Bakari Ponda, and Loy “Babu” Loishoki. Funding was provided by NSF Doctoral Dissertation Improvement Grants (BCS-0925901, BCS-0925690), the Leakey Foundation, Primate Conservation Inc., Primate Action Fund, Margot Marsh Biodiversity Foundation, Sigma XI, Idea Wild, and Stony Brook University’s Interdepartmental Doctoral Program in Anthropological Sciences Research Fund.

### SUPPLEMENTARY MATERIALS AND METHODS

***Traditional, frequentist meta-analyses***

To assess the robustness of our Bayesian meta-analyses, we constructed complementary frequentist models using the ‘metafor’ package (2) in ‘R.’ The results of these models were largely concordant with those provided in the Main Text. However, these models differ in their estimation of random effects variances, and thus their outputs somewhat differed. In particular, frequentist meta-analyses are prone to underestimating between-study variance, resulting in zero estimates for random effects parameters (3, 4). By contrast, Bayesian meta-analyses are prone to overestimating between-study variance, which often produces more conservative estimates (5). Thus, the patterns documented in the Main Text reflect a more conservative approach, and random effects variances were occasionally estimated at zero when using traditional, frequentist meta-analyses. We summarize these key differences below.

Patterns regarding female-female relationships were broadly equivalent to those described in the Main Text. Females in the single-level societies formed stronger relationships with kin, and these patterns were stronger in the cliquish societies than in the cohesive societies (**Figure S2a**). Likewise, females in the multi-level geladas showed very strong kin biases, while females in multi-level Guinea baboons showed weaker kin biases. Rank similarity effects were similar to those detected using Bayesian approaches. Rank effects were present in the cohesive groups, stronger in the cliquish groups, and absent in the multi-level societies (**Figure S2b**). Again, shared male effects were only present in the multi-level societies (**Figure S2c**).

Patterns regarding female-male relationships, however, showed more differences across the two methods. Using frequentist approaches, male rank effects were weak-but-detectable in the cohesive and cliquish societies but considerably stronger in the multi-level societies (**Figure S2d**). Female rank effects were weakly present in the cliquish papionins (**Figure S2e**). Lastly, rank interaction effects were weak but present in only the cliquish societies (**Figure S2f**). Further inspection of the data reveals that these different outputs may result because Bayesian methods are more skeptical of effects when species differences are present. In particular, male rank effects were stronger in mandrills and mangabeys than in the single-level baboons. Likewise, female rank and rank interaction effects were often salient in chacma baboons and mandrills but not in Kinda baboons. Thus, pooling effect sizes across these taxa led to negligible effects when using Bayesian methods, which attribute a greater proportion of variance to random effects.

***Imputation methods***

Due to differences in the depth of pedigree and known genetic relationships across the study populations, kinship information was not available for all female-female dyads (13.3% of female-female dyads). Likewise, rank information was missing for a small proportion of dyads (3.1% of female-female dyads, 6.6% of all female-male dyads, as males who briefly passed through social groups were not always assigned ranks). Using only dyads with known information, however, could generate incorrect or biased inferences, particularly when estimating kinship effects. For instance, in populations with shallower pedigrees, many mother-offspring and maternal sister dyads are known but fewer more distant kin dyads can be determined from observation alone. As a result, the mean strength of grooming relationships among kin could be artificially inflated in these populations. To ameliorate these concerns, we used the ‘mice’ package (1) in ‘R’ to impute missing relatedness and rank data using a classification and regression trees approach, which accommodates both continuous and categorical data and implicitly discerns interaction effects across included variables (i.e., population-specific effects, etc.).

***Kinship data.*** To impute missing kinship data in female-female dyads, we calculated lifetime grooming indices for each dyad within the entire dataset (n = 11363 unique dyads). Specifically, we calculated the undirected grooming rate for each dyad-year; divided this value by the mean grooming rate across all dyads within the respective group-year, thereby standardizing this measure;

and averaged these grooming indices across all of the years of the dataset to create a lifetime relationship strength metric. Thus, each dyad was distilled to a single observation to ensure that imputed binary kinship measures (yes/no) were equivalent across all years the dyad was co-resident. We used the following covariates as informative criteria for imputation: (i) study species; (ii) study population; (iii) edge definition (i.e., proportion vs. rates); (iv) the species's dispersal pattern, as kin dyads should be more prevalent in female-philopatric groups; (vi) the dyad's mean absolute rank difference, as related dyads tend to also be close in rank; and (vii) the dyadic grooming index described above, square-root transformed to reduce the impact of extreme values.

To verify the predictive accuracy of these imputation procedures, we removed 30% of all confirmed kin and non-kin dyads, repeated the same imputation procedures, and quantified the proportion of relationships that were correctly recovered by the imputation process across 100 replicates. Notably, classification and regression tree imputation recovered 'true positive' and 'true negative' kin relationships more often than expected by chance alone (**Figure S5**). Although the 'true positive' rate was still low (<40% on average, as false negatives were prevalent), these errors did not systematically bias our results. 'True positive' kin dyads formed very strong grooming relationships, and true negative non-kin dyads formed extremely weak grooming relationships. However, 'false positive' and 'false negative' grooming relationships were broadly similar in their strength and were equally prevalent (**Figure S6a**). Given these equivalencies, kin bias estimates generated using dyads of known kinship ($\beta_{kinship.true}$ = 0.98, 89% CI = [0.83, 1.13]) were broadly consistent with and fully overlapped estimates generated using imputed data ($\beta_{kinship.imputed}$ = 0.87, 89% CI = [0.73, 1.01]; **Figure S6b,c**).

***Rank data.*** To impute missing rank data, we aggregated grooming data from each node within the annual grooming networks. We then imputed missing proportional rank values using the following covariates: (i) study species; (ii) study population; (iii) edge definition (i.e., proportion vs. rates); (iv) grooming out-strength (i.e., the sum of the grooming *given*); and (v) grooming in-strength (i.e., the sum of grooming *received*). Prior to imputation, these two grooming network metrics were z-scored relative to other observations from that respective group-year. We imputed data on males and females separately, as the sexes likely follow different rank patterns. Given this set of informative covariates, imputed rank values were generated based on species- and population-specific relationships between dominance rank and grooming interactions.

To verify the accuracy of these imputation procedures, we removed known rank values for 30% of all female and male nodes, repeated these imputation methods, and quantified the correlation between the observed rank values and those identified across 100 imputed datasets. For both females and males, imputation methods produced stronger Pearson correlations than expected due to random chance alone (Kolmogorov-Smirnov test, female nodes: $D = 0.52$, $P < 0.001$; male nodes: $D = 1$, $P < 0.001$), although predictive accuracy was much higher for males (median Pearson's $r = 0.36$) than for females (median Pearson's $r = 0.078$). Despite these broadly low correlations between observed and imputed data, models using only imputed data recovered the observed pattern, in which high-ranking individuals tended to receive more grooming than their low-ranking counterparts. In other words, analyses based only on dyads with known ranks did not capture different grooming patterns than analyses including these imputed dyads. Given this consistent relationship and the small fraction of missing information, including imputed data did not alter the relationship between rank on grooming behavior. Nevertheless, we opted to use imputation methods and retain dyads with missing rank information, as this allowed us to preserve statistical power to assess other effects of interest (e.g., shared male effects) when quantifying grooming interactions in small social groups.

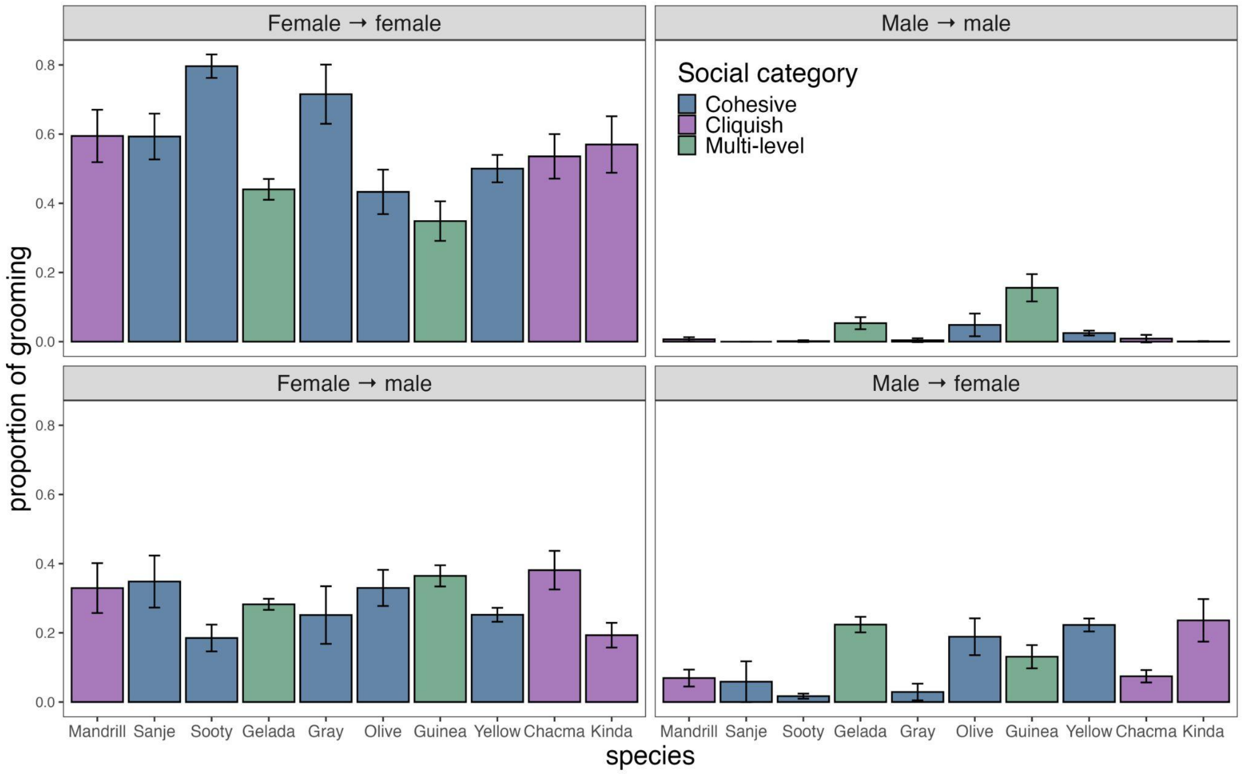


**Figure S1. Proportion of group-level grooming budgets, divided by the sex-specific direction of grooming.** Female-female grooming comprises the bulk of grooming for nearly all papionin species. Concomitantly, the relative importance of female-to-male and male-to-female grooming varies widely across the papionins. Male-male grooming was infrequent in most papionin species. Two exceptions to this are Guinea baboons and geladas, in which co-resident leader males and follower males regularly engage in grooming. A third exception is hamadryas baboons, in which solitary and follower males engage in frequent male-male grooming. However, the hamadryas data set used for this study did not include these males, who are part of clans and bands but are not part of one-male units. Given the lack of data on these males, we did not include hamadryas in the above comparison. Error bars indicate means ± 2 SE for each measurement, pooled across the 20 posterior networks generated using 'bisonR'. Species are colored according to their social category, as described in the Main Text.

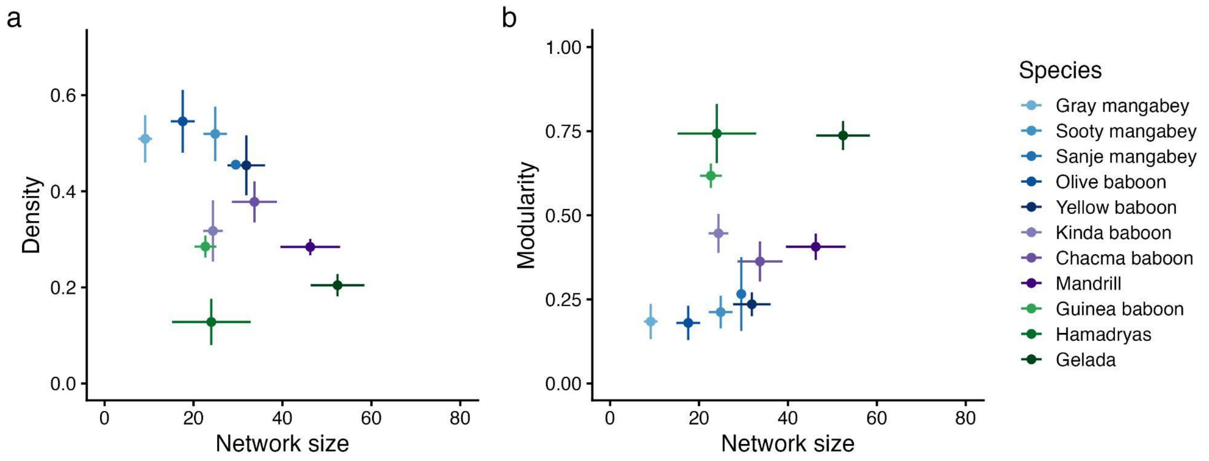


**Figure S2. Variation in network metrics and network size across species.** Species differences in grooming network (a) density and (b) modularity persisted across network sizes. Points and vertical error bars indicate mean values ± 2 SE (pooled across the 20 posterior networks) for each respective network metric, while horizontal error bars indicate mean group sizes ± 2 SE.

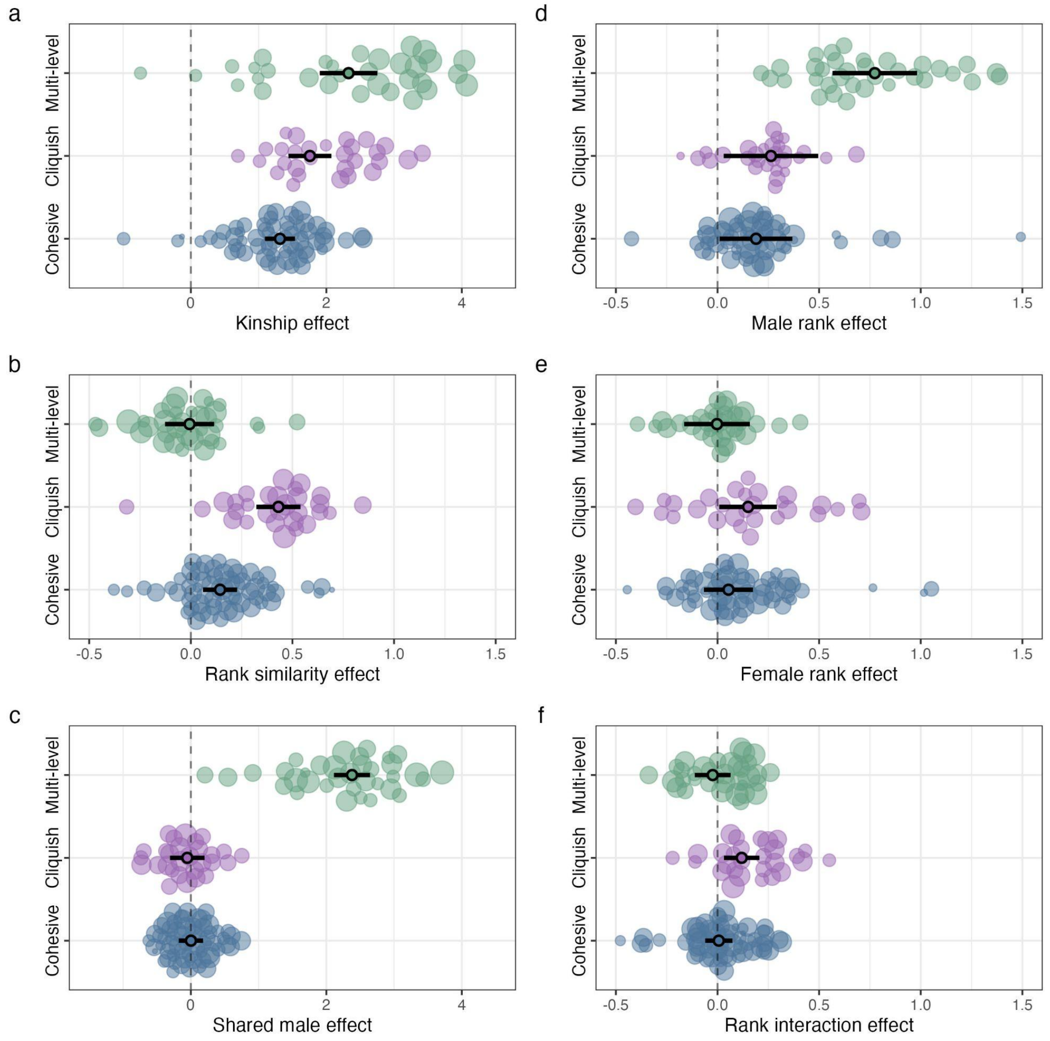


**Figure S3. Results of frequentist meta-analyses.** (a). Kinship was a salient predictor of female-female grooming relationships, although this pattern varied across social systems and female dispersal patterns. (b) Rank similarity effects were generally positive, indicating that closely-ranked females groomed more often. However, these tendencies were strongest in cliquish societies and absent in multi-level societies. (c) Females that shared the same top male partners formed stronger grooming relationships in the multi-level groups but not in the single-level groups. (d) High-ranking and reproductively dominant males received more grooming from females, and this tendency was strongest in the multi-level societies. (e) High-ranking females did not typically form stronger between-sex relationships. (f) Rank interaction effects were present only in the cliquish societies. This pattern differs from that detected from the Bayesian models. Each point indicates a standardized effect size for the corresponding predictor. Black points and error bars indicate means and 95% confidence intervals.The size of each point is scaled to 1 divided by the effect size's standard error. Thus, effects with greater precision are represented by larger points.

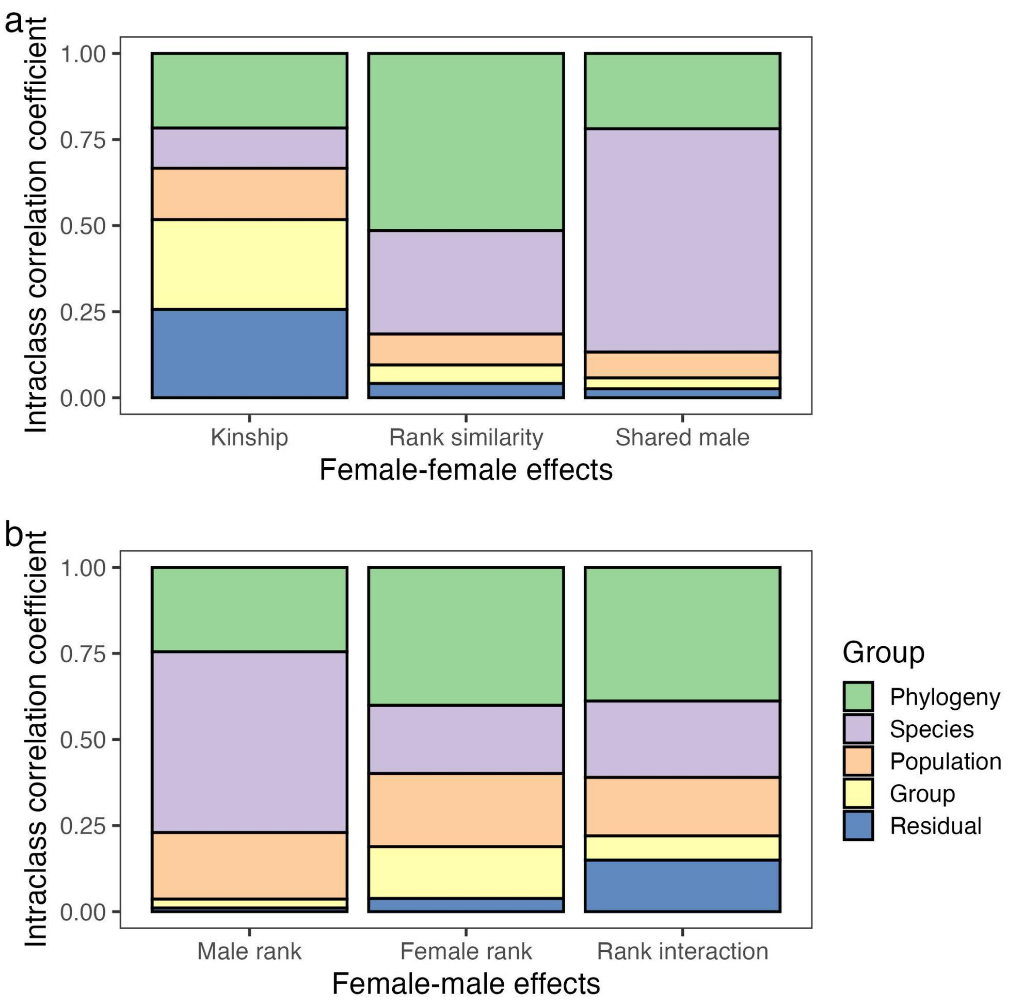


**Figure S4. Variance decomposition for (a) female-female and (b) female-male dyadic regression models.** Intra-class correlations for random effects across each meta-analysis. These values indicate the proportion of variance explained by each of these random effects.

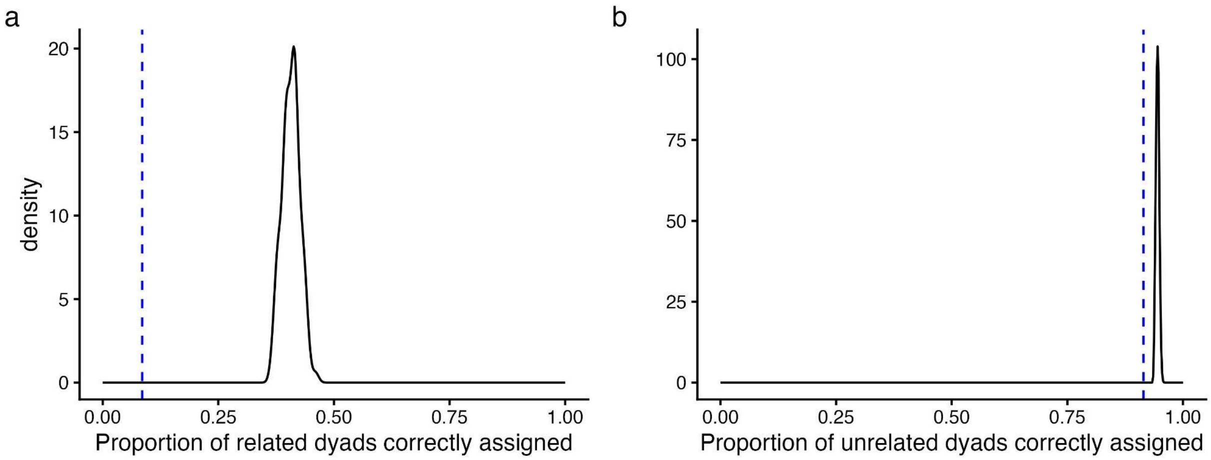


**Figure S5**. **Imputation correctly assigned related and unrelated dyads more often than expected by chance.** The black density peaks indicate the proportion of dyads that were correctly assigned as (a) related and (b) unrelated across 100 imputations. The dashed blue line indicates the proportion of dyads that would be assigned correctly using random resampling.

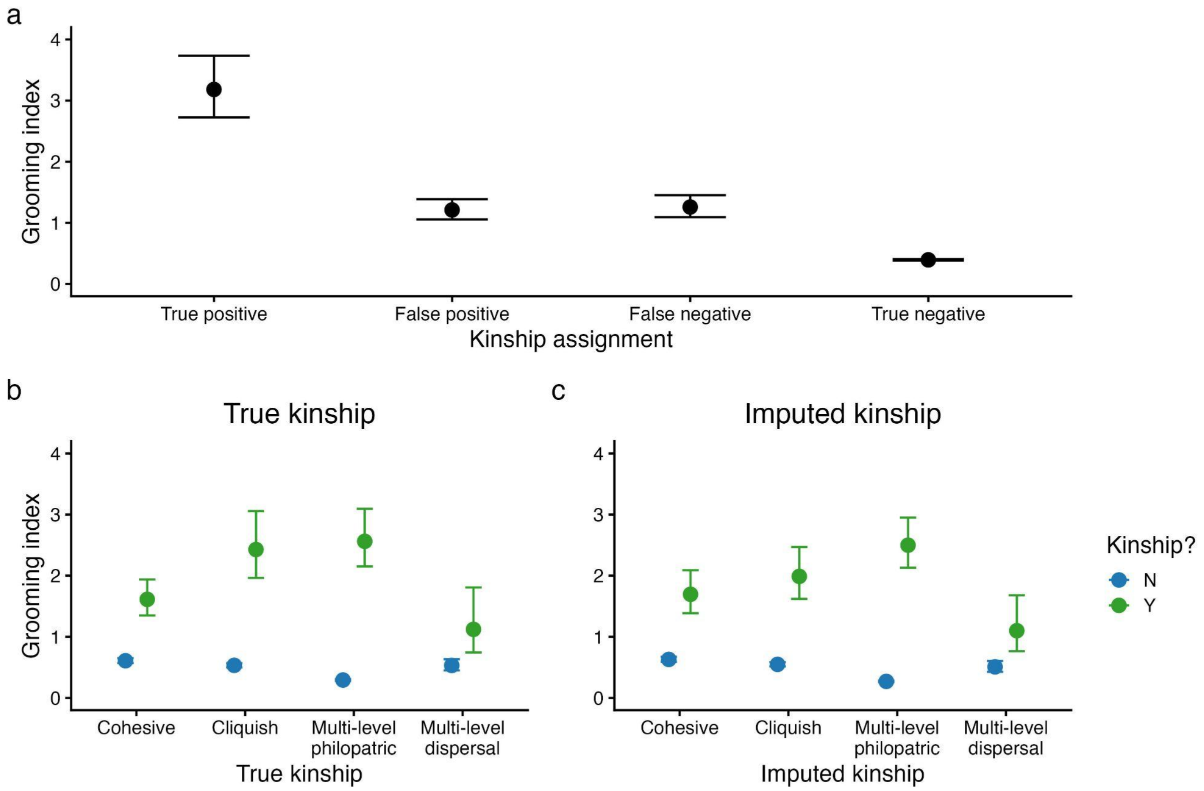


**Figure S6**. **Grooming relationship strengths varied across kinship assignments.** True positive kin dyads formed the strongest grooming relationships, while true negative non-kin dyads formed the weakest grooming relationships. False positive and false negative dyads were largely equivalent in their grooming strengths and in their prevalence. Thus, imputation broadly recovers comparable patterns of kin bias in grooming relationships.

**Table S1. Demographic characteristics of sampled populations and social groups.** The average number of sampled adults and subadults per social group across the study populations. Data from single-level societies were collected at the group-level, while data from Guinea baboons were collected at the party-level, hamadryas baboons at the clan-level, and geladas at the band-level. For the single-level groups and Guinea baboon parties, these values note the total average number of subadult and adult individuals, excluding on occasion individuals who were poorly sampled or reached maturity too far into the group-year. For gelada bands and hamadryas baboon clans, however, these values note the average number of individuals from individually recognized and sampled one-male units and not the size of the entire multi-level society. Thus, these numbers represent an undercount of total group size, as they exclude unsampled units and, in hamadryas baboons, solitary males. The number of group-years of data provided by each study is also noted.

| Species | Population | $N_{females}$ | $N_{males}$ | $N_{groups}$ | $N_{group-years}$ | Obs. Effort*** | Study Years | Citation |
|---|---|---|---|---|---|---|---|---|
| *Cercocebus sanjei* | Udzungwa, Tanzania | 21.5 | 8.0 | 1 | 2 | 54.4-56.8 | 2009-2010 | (6) |
| *Cercocebus atys atys* | Taï Forest, Côte d'Ivoire | 20.1 | 4.8 | 1 | 8 | 22.6-121.8 | 2014-2021 | (7) |
| *Lophocebus albigena* | Kibale, Uganda | 6.6 | 2.5 | 4 | 15 | 2.7-14.6 | 2010-2013 | (8) |
| *Papio anubis* | Laikipia, Kenya | 13.4 | 6.9 | 3 | 9 | 11.1-23.2 | 2013-2017 | (9) |
| *Papio anubis* | Gashaka, Nigeria | 9.8 | 5.7 | 2 | 12 | 14.0-174.9 | 2009-2016 | (10) |
| *Papio cynocephalus** | Amboseli, Kenya | 19.4 | 12.4 | 5 | 25 | 2.5-12.4 | 2003-2007 | (11) |
| *Papio kindae* | Kasanka, Zambia | 19.9 | 4.5 | 1 | 8 | 4.2-16.3 | 2011-2018 | (12) |
| *Papio ursinus* | Moremi, Botswana | 29.0 | 13.2 | 1 | 9 | 2.3-13.8 | **1992-2007 | (13) |
| *Papio ursinus* | Tsaobis, Namibia | 16.4 | 7.8 | 2 | 8 | 16.0-67.7 | **2006-2014 | (14) |
| *Mandrillus sphinx* | Lékédi, Gabon | 35.0 | 11.3 | 1 | 4 | 7.9-13.3 | 2013-2016 | (15) |

| | | | | | | | | |
|---|---|---|---|---|---|---|---|---|
| *Papio papio* | Simenti, Senegal | 12.3 | 10.3 | 3 | 12 | 6.9-13.8 | 2016-2019 | (16) |
| *Papio hamadryas* | Filoha, Ethiopia | 14.7 | 9.3 | 2 | 3 | 5.8-14.2 | 1997-1998 | (17) |
| *Theropithecus gelada* | Simien Mountains, Ethiopia | 38.5 | 14.0 | 2 | 20 | 1.8-17.3 | 2009-2019 | (18) |

*The baboon population at Amboseli consists of *P. cynocephalus/anubis* hybrids (19). Despite a long history of introgression, *P. cynocephalus* still makes up the bulk of the population's genetic background. **Indicates studies with gaps between sampling periods included within the CAPS database. ***Average observation hours per capita across all group-years in the dataset.

**Table S2. Results of the Bayesian models focused on the drivers of grooming network structure.** Parameter estimates for models assessing how network size, social structure categories, and sampling effort are linked with social network structure (n = 135 networks). Phylogeny, species, population, and group were included as random effects in all models.

| Model | Fixed effect | Mean | Est. Error | 89% CI LL | 89% CI UL |
|---|---|---|---|---|---|
| ***Density*** | Intercept | -0.09 | 0.31 | -0.55 | 0.44 |
| | $\beta_{NetworkSize}$ | -0.55 | 0.08 | -0.69 | -0.42 |
| | $\beta_{Cliquish}$ | -0.23 | 0.42 | -0.88 | 0.43 |
| | $\beta_{Multi\text{-}level}$ | -0.89 | 0.42 | -1.53 | -0.23 |
| ***Modularity*** | Intercept | -1.15 | 0.22 | -1.49 | -0.82 |
| | $\beta_{NetworkSize}$ | 0.30 | 0.10 | 0.15 | 0.48 |
| | $\beta_{Cliquish}$ | 1.63 | 0.29 | 0.16 | 1.05 |
| | $\beta_{Multi\text{-}level}$ | 1.77 | 0.28 | 1.32 | 2.19 |

**Table S3. Description of female-female and female-male dyadic regression models constructed for each group-year within the dataset**. Description of the dataset and dyadic modelling approach for female-female and female-male dyadic regressions. Separate dyadic regressions were conducted for each group-year in the dataset (n = 126 for female-female dyads, n = 124 for female-male dyads, as we removed group-years where group size limited our ability to jointly estimate these effects).

| Model type | $N_{dyad\text{-}years}$ | Fixed effects | Random effects |
|---|---|---|---|
| Female-female | 35495<br>(6-1705 per group-year) | Kinship (yes/no)*<br>Rank similarity<br>Shared male (yes/no) | mm(ID1, ID2) |
| Female-male | 28218<br>(15-1086 per group-year) | Male rank<br>Female rank<br>Rank interaction | mm(ID1, ID2) |

*Kinship information was not available for hamadryas baboons. Thus, we removed this covariate from the three models focused on this species.

**Table S4. Results from the Bayesian meta-analyses focused on the determinants of the strength of kin biases, rank effects, and shared male patterns.** Parameter estimates for Bayesian models examining the effects of social system category and group size on female-female and female-male dyadic effects.

| Outcome | Moderator | Mean effect size | 89% CI LL | 89% CI UL |
|---|---|---|---|---|
| ***Female-female relationships*** **(n = 126*)** | | | | |
| Kinship* | $\beta_{Cohesive}$ | 1.34 | 1.01 | 1.65 |
| | $\beta_{Cliquish}$ | 1.82 | 1.41 | 2.21 |
| | $\beta_{Multi\text{-}level}$ | 2.34 | 1.69 | 2.93 |
| | $\beta_{Female\ dispersal}$ | -1.14 | -1.95 | -0.27 |
| | $\beta_{Network\ size}$ | 0.46 | 0.31 | 0.63 |
| Rank similarity | $\beta_{Cohesive}$ | 0.15 | 0.01 | 0.28 |
| | $\beta_{Cliquish}$ | 0.44 | 0.27 | 0.60 |
| | $\beta_{Multi\text{-}level}$ | -0.04 | -0.29 | 0.23 |
| | $\beta_{Dispersal}$ | 0.12 | -0.21 | 0.45 |
| | $\beta_{Network\ size}$ | -0.01 | -0.06 | 0.04 |
| Shared male | $\beta_{Cohesive}$ | 0.01 | -0.23 | 0.25 |
| | $\beta_{Cliquish}$ | -0.09 | -0.38 | 0.20 |
| | $\beta_{Multi\text{-}level}$ | 2.15 | 1.62 | 2.63 |
| | $\beta_{Dispersal}$ | 0.49 | -0.14 | 1.17 |
| | $\beta_{Network\ size}$ | 0.03 | -0.11 | 0.17 |
| ***Female-male relationships*** **(n = 124)** | | | | |
| Male rank | $\beta_{Cohesive}$ | 0.19 | 0.01 | 0.37 |
| | $\beta_{Cliquish}$ | 0.26 | 0.03 | 0.50 |
| | $\beta_{Multi\text{-}level}$ | 0.78 | 0.57 | 0.98 |
| | $\beta_{Network\ size}$ | -0.01 | -0.08 | 0.06 |
| Female rank | $\beta_{Cohesive}$ | 0.05 | -0.07 | 0.17 |
| | $\beta_{Cliquish}$ | 0.15 | 0.01 | 0.29 |
| | $\beta_{Multi\text{-}level}$ | -0.00 | -0.16 | 0.16 |
| | $\beta_{Network\ size}$ | 0.03 | -0.03 | 0.08 |
| Rank interaction | $\beta_{Cohesive}$ | 0.00 | -0.06 | 0.07 |
| | $\beta_{Cliquish}$ | 0.12 | 0.03 | 0.21 |
| | $\beta_{Multi\text{-}level}$ | -0.02 | -0.11 | 0.06 |
| | $\beta_{Network\ size}$ | 0.04 | -0.00 | 0.08 |

*Kinship information was not available for hamadryas baboons (n = 3 group-years). Thus, the sample size for kinship effects was 123 group-years.

**Table S5. Results of the traditional, frequentist meta-analyses focused on the determinants of the strength of kin biases, rank effects, and shared male patterns.** Parameter estimates for frequentist meta-analytic models examining the effects of social system category and group size on female-female and female-male dyadic effects. Here, 95% CI denotes the confidence intervals for each respective effect.

| Outcome | Moderator | Mean effect size | 95% CI LL | 95% CI UL |
|---|---|---|---|---|
| ***Female-female relationships*** **(n = 126*)** | | | | |
| Kinship* | $\beta_{Cohesive}$ | 1.43 | 1.23 | 1.63 |
| | $\beta_{Cliquish}$ | 1.87 | 1.56 | 2.19 |
| | $\beta_{Multi\text{-}level}$ | 2.44 | 1.97 | 2.92 |
| | $\beta_{Female\ dispersal}$ | -1.22 | -1.93 | -0.52 |
| | $\beta_{Network\ size}$ | 0.49 | 0.31 | 0.66 |
| Rank similarity | $\beta_{Cohesive}$ | 0.15 | 0.08 | 0.22 |
| | $\beta_{Cliquish}$ | 0.43 | 0.34 | 0.53 |
| | $\beta_{Multi\text{-}level}$ | -0.09 | -0.24 | 0.07 |
| | $\beta_{Dispersal}$ | 0.16 | -0.07 | 0.39 |
| | $\beta_{Network\ size}$ | 0.01 | -0.05 | 0.07 |
| Shared male | $\beta_{Cohesive}$ | 0.02 | -0.16 | 0.21 |
| | $\beta_{Cliquish}$ | -0.07 | -0.33 | 0.19 |
| | $\beta_{Multi\text{-}level}$ | 2.12 | 1.71 | 2.53 |
| | $\beta_{Dispersal}$ | 0.50 | -0.09 | 1.10 |
| | $\beta_{Network\ size}$ | 0.03 | -0.13 | 0.19 |
| ***Female-male relationships*** **(n = 124)** | | | | |
| Male rank | $\beta_{Cohesive}$ | 0.26 | 0.02 | 0.49 |
| | $\beta_{Cliquish}$ | 0.23 | -0.08 | 0.53 |
| | $\beta_{Multi\text{-}level}$ | 0.64 | 0.34 | 0.94 |
| | $\beta_{Network\ size}$ | 0.04 | -0.05 | 0.13 |
| Female rank | $\beta_{Cohesive}$ | 0.06 | -0.08 | 0.19 |
| | $\beta_{Cliquish}$ | 0.12 | -0.06 | 0.30 |
| | $\beta_{Multi\text{-}level}$ | -0.06 | -0.25 | 0.12 |
| | $\beta_{Network\ size}$ | 0.05 | -0.03 | 0.12 |
| Rank interaction | $\beta_{Cohesive}$ | 0.04 | -0.04 | 0.12 |
| | $\beta_{Cliquish}$ | 0.16 | 0.04 | 0.27 |
| | $\beta_{Multi\text{-}level}$ | 0.01 | -0.11 | 0.13 |
| | $\beta_{Network\ size}$ | 0.01 | -0.03 | 0.06 |

*Kinship information was not available for hamadryas baboons (n = 3 group-years). Thus, the sample size for kinship effects was 123 group-years.
